# Deficiency in Translesion DNA Polymerase ζ Induces an Innate Immune Response

**DOI:** 10.1101/2020.03.02.972513

**Authors:** Sara K. Martin, Junya Tomida, Richard D. Wood

**Affiliations:** Department of Epigenetics & Molecular Carcinogenesis, The University of Texas MD Anderson Cancer Center, Smithville, Texas, USA; The University of Texas MD Anderson Cancer Center UT Health Graduate School of Biomedical Sciences, Smithville, Texas, USA; Department of Biological Sciences, University of North Carolina, Charlotte, USA

## Abstract

DNA polymerase pol ζ is regarded as a specialized DNA polymerase for bypass of DNA lesions. In mammalian cells, pol ζ also contributes to genomic stability during normal DNA replication. Disruption of *Rev3l* (the catalytic subunit of pol ζ) is toxic to cells and mice, with increased constitutive chromosome damage, including micronuclei. As the cellular manifestations of this genomic stress have remained unexplored, we measured genome-wide transcriptional changes by RNA-seq in pol ζ-defective cells. Expression of 1117 transcripts was altered by 4-fold or more in *Rev3l* knockout mouse embryonic fibroblasts (MEFs), with a pattern showing an induction of an innate immune response. We validated the increased expression of known interferon-stimulated genes (ISG) at the mRNA and protein levels. We found that the cGAS-STING axis, which senses cytosolic DNA, drives ISG expression in *Rev3l* knockout MEFs. These results reveal a new genome protective function of pol ζ and indicate that inhibition of pol ζ may be therapeutically useful by simultaneously increasing sensitivity to genotoxins and inducing a cytotoxic innate immune response.

## INTRODUCTION

Mammalian genomes encode an array of translesion DNA polymerases, which provide a diverse tool kit to tolerate assorted genomic lesions [1]. While most translesion DNA polymerases are required for cells to survive various exogenous genotoxic assaults, they are not essential for mammalian development or unchallenged cellular survival [1]. An exception is DNA polymerase ζ (pol ζ) whose catalytic subunit is encoded by the *Rev3l* gene [2]. Germline disruption of *Rev3l* results in embryonic lethality in mice [3–5]. The indispensable nature of pol ζ likely stems from inadequately understood genome protective functions. Disruption of *Rev3l* in B cells or keratinocytes in mice leads to acute genomic stress in the target tissues [6–10]. Primary mouse embryonic fibroblasts (MEFs) rapidly accumulate chromosome breaks at the first metaphase following *Rev3l* disruption [11]. Loss of pol ζ in primary MEFs cripples cell proliferation, with the cells failing to replicate past roughly two rounds of cell division following *Rev3l* disruption [11]. The dramatic growth suppression in *Rev3l* knockout MEFs appears to be ameliorated by blunting DNA damage quality control responses, for example by p53 deletion [12] or T-antigen expression [11] (which inhibits p53 and other targets). However, p53 deletion utterly fails to rescue embryonic lethality of *Rev3l* disruption in mice [12–14]. This implies that there are additional, p53-independent, growth suppressive responses to the genomic strain caused by *Rev3l* disruption. We set out to clarify the consequences of the constitutive genomic stress induced by the loss of pol ζ in mammalian cells. Starting with an unbiased transcriptome wide approach, we discovered that loss of pol ζ induces constitutive activation of an innate immune response. Further we found that the cytosolic nucleic acid sensor, cGAS, and its downstream signalling partner STING drive this innate immune response in absence of functional pol ζ. Given that the cGAS-STING axis can inhibit cell growth, this gives a potential explanation for why loss pol ζ function is remarkably toxic to proliferating cells.

## RESULTS & DISCUSSION

### A shortened REV3L construct rescues known phenotypes of pol ζ disruption

In order to dissect the consequences of the poorly resolved genome protection function of pol ζ, we set up a complementation system using T-antigen immortalized mouse embryonic fibroblasts (MEFs) [11] either with a pol ζ proficient background, *Rev3l* heterozygous (HET) background, or a pol ζ deficient background, *Rev3l* knockout (KO). A biochemically active shortened human REV3L construct, TR4-2 [15] (Fig 1A) with an N-terminal Flag-HA tag was stably expressed in *Rev3l* KO MEFs (Fig 1B, Fig S1A). In order to test that our complementation system was in fact addressing both external and endogenous genome protective functions of pol ζ, we tested both cellular sensitivity to cisplatin and formation of micronuclei in unchallenged cells, one marker of genomic instability. As expected *Rev3l* KO + empty vector MEFs were hypersensitive to cisplatin relative to the *Rev3l* HET MEFs + empty vector (Fig 1C, Fig S1B). Stable expression of TR4-2 in *Rev3l* KO MEFs reversed the hypersensitivity to cisplatin (Fig 1C, Fig S1B). This indicates that in this context TR4-2 can restore REV3L’s function in DNA damage tolerance. Consistent with large scale genomic stress, ~15-23% of unchallenged *Rev3l* KO MEFs had at least one micronucleus, relative to *Rev3l* HET MEFs which had a micronucleus frequency of ~1-3% (**Fig 1D**). Stable TR4-2 expression in *Rev3l* KO MEFs restored micronucleus frequency to near-normal, showing that TR4-2 can restore some *Rev3l* genome protective functions. We used this isogenic system as a tool to probe the unknown consequence of the loss of pol ζ.

**Figure 1:**
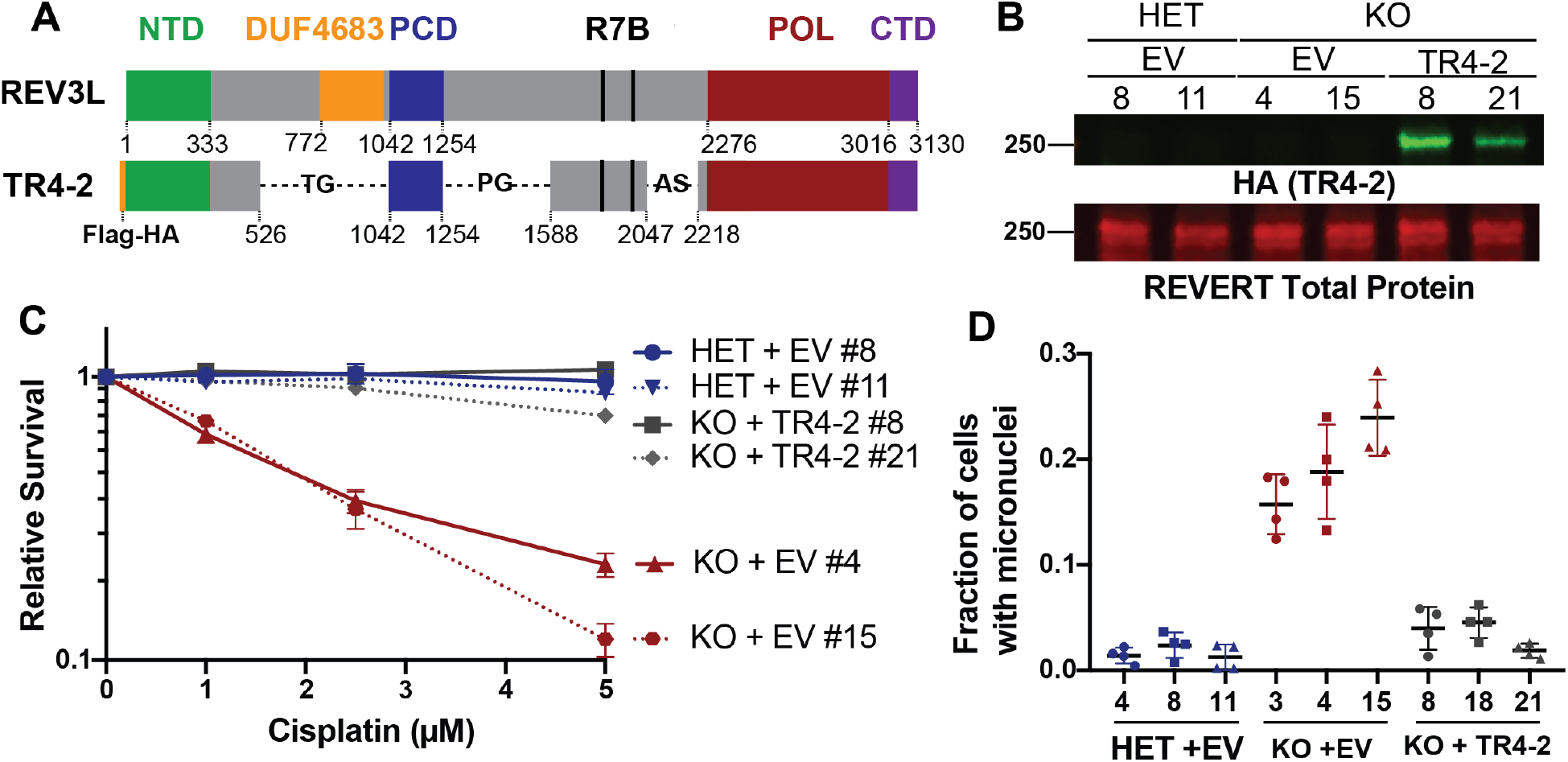
Shortened REV3L construct rescues phenotypes of pol ζ disruption. A) Schematic of full-length human REV3L and human REV3L construct, TR4-2. TR4-2 retains most conserved domains and binding sites of REV3L including the regions that coordinate interactions with the accessory subunits of pol ζ, the C-terminal domain (CTD, purple) the two REV7 binding sites (R7B, black), and a positively charged domain (PCD) of uncertain function. The B-family catalytic core is formed by folding of the N-terminal domain (NTD, green) and the polymerase domain (POL, red) [2]. B) Immunoblot with HA antibody showing stable expression of TR4-2 with an N-terminal Flag-HA tag in *Rev3l* KO MEF clones. Full blots are shown in Fig S2. C) Stable expression of TR4-2 in *Rev3l* KO MEF clones reverses hypersensitivity to cisplatin. MEFs were exposed to the indicated cisplatin concentrations for 48 h and relative survival was quantified with the ATPlite assay. D) Stable expression of TR4-2 decreases micronuclei formation in unchallenged *Rev3l* KO MEF clones.

### Loss of polymerase ζ alters the transcriptome

In order to uncover the type of stress occurring in cells lacking pol ζ, we performed genome-wide mRNA sequencing on a controlled set of immortalized clones: *Rev3l* HET + empty vector, *Rev3l* KO + empty vector and *Rev3l* KO + TR4-2. To focus on major changes, we set a strict threshold (> |2| log_2_ fold change and FDR < 0.05). Expression analysis of 17,346 mapped transcripts revealed that 1117 transcripts were either upregulated or downregulated in the *Rev3l* KO + empty vector relative to the *Rev3l* HET + empty vector MEFs (Fig 2A). The majority (~68%) were upregulated (Fig 2A). These upregulated or downregulated genes displayed no statistically significant enrichment or depletion for DNA replication or canonical DNA damage sensing pathways. This is not completely unexpected given that p53 promotes much of the transcriptional response to DNA damage, while our MEFs have inactivated p53 due to large T-antigen immortalization. In these immortalized high passage cells, we are likely to observe stable transcriptional alterations, rather than an acute response.

**Figure 2:**
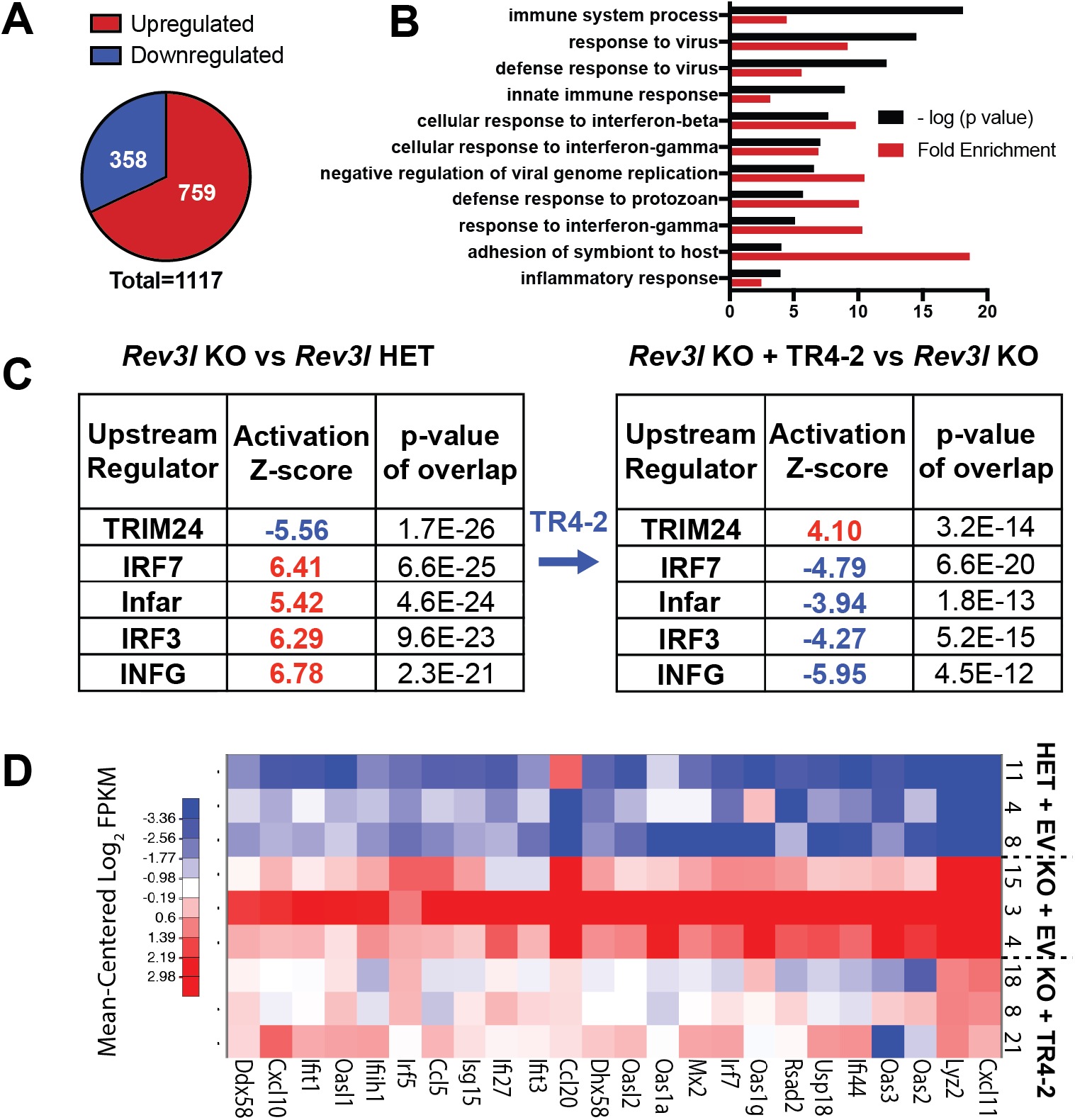
Cells deficient in DNA polymerase ζ have an altered transcriptome. A) Differentially expressed genes in *Rev3l* KO + empty vector relative to *Rev3l* HET + empty vector using the threshold of a log_2_ fold change of > |2| and a false discovery rate of < 0.05. B) Top 10 GO (Gene Ontology) terms reveal enrichment of immune system related genes in upregulated genes in *Rev3l* KO MEFs. C) Upstream regulator analysis reveals the data set is consistent with predicted activation of positive regulators of an interferon response in *Rev3l* KO MEFs. D) Heatmap of a set of 25 known interferon-stimulated genes.

Instead the upregulated genes displayed an enrichment in immune system-related pathways, as revealed by gene ontology analysis (Fig 2B). Upstream regulator analysis was used to analyze all differentially expressed genes. This revealed that the alterations in the transcriptome are consistent with activation of positive regulators of the interferon response, including key transcription factors in this pathway, IRF3 and IRF7 (Fig 2C). Importantly the predicted activation of IRF3 and IRF7 was reversed by expression of the *Rev3l* TR4-2 cDNA, showing that these results stem from a function of *Rev3l.* The same trends were also observed by applying a substantially lower threshold (log_2_ FC > |0.5|) for differentially expressed genes. This increased the dataset to 2071 differentially expressed transcripts. Pathway analysis showed a negative correlation with predicted TRIM24 activation. TRIM24 suppresses interferon-stimulated gene expression [16], confirming that our data is consistent with expression of interferon stimulated genes.

To explore whether our data set is in fact consistent with an interferon response, we analyzed a curated set of 25 known interferon stimulated genes and observed increased expression in the *Rev3l* KO relative to the *Rev3l* HET MEFs, which was partially reduced by TR4-2 stable expression (Fig 2D). Together our results reveal that disruption of pol ζ promotes induction of interferon-stimulated genes.

### Disruption of *Rev3l* induces interferon-stimulated gene expression driven by the cGAS-STING axis

Given that our complemented cell lines were generated using lentivirus constructs and grown continually under selection, we moved to validate the results in the parental *Rev3l* KO and *Rev3l* HET MEF cell lines and one additional set of cell lines to limit extraneous variables. We confirmed an increase in mRNA expression of specific interferon stimulated genes in the *Rev3l* KO MEFs relative to the control cell lines, including key chemokines (Fig 3A). In order to extend these findings to the protein level, we examined interferon stimulated gene products by immunoblotting. Corresponding to an increase in mRNA levels, we also observed an increase in protein levels of known interferon stimulated genes, MDA5 (encoded by the *IFIH1* gene), ISG15, and viperin (encoded by the *RSAD2* gene) (Fig 3B). Together these data indicate that an interferon branch of the innate immune system may be activated due to disruption of pol ζ function. Since it seems unlikely that pol ζ plays a direct role in transcriptional regulation, the next obvious question is how and why loss of pol ζ induces the expression of interferon stimulated genes.

**Figure 3:**
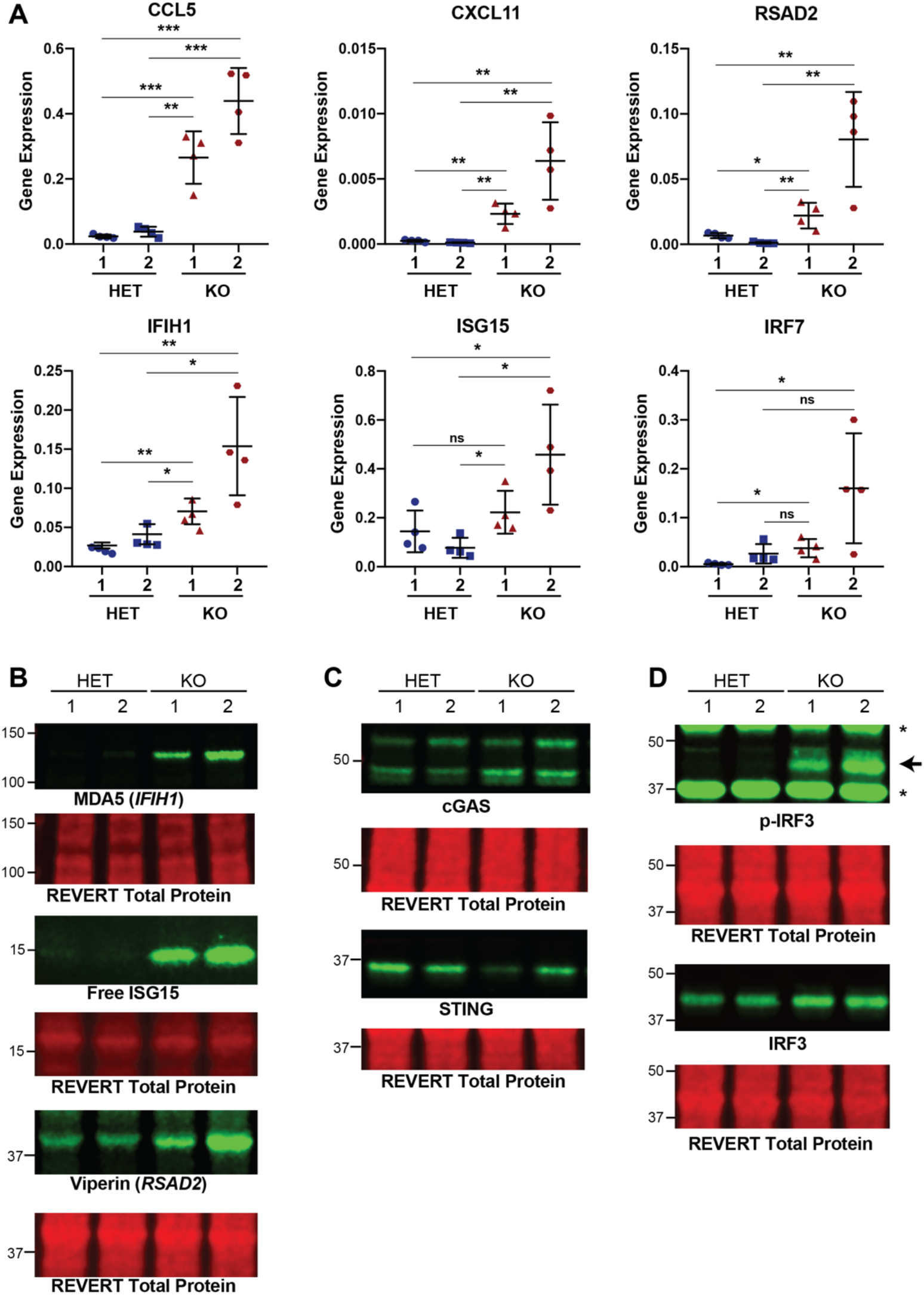
Disruption of *Rev3l* results in increased expression of interferon stimulated genes and proteins. A) Gene expression (2^−ΔCt^) of selected interferon stimulated genes normalized to HPRT detected by qRT-PCR. Error bars represent standard deviation. Unpaired student t-test, * = p < 0.05, ** = p < 0.01 and *** = p < 0.001. B) Immunoblots showing increased protein levels of interferon stimulated genes, MDA5 and ISG15. C) Immunoblot showing presence of components of the innate immune system, cGAS and STING, in MEFs, with reduced STING in pol ζ knockout cells. D) Enhanced phosphorylation of S888 of IRF3 (corresponding to S396 in humans) in *Rev3l* KO MEFs. Full blots are shown in Fig S3.

The major consequence of pol ζ disruption in unchallenged mammalian cells is increased genomic instability as evidenced by multiple markers including γ-H2AX foci, chromosome fragmentation and aberrations, and micronuclei [10–12,17] (Fig 1D). Therefore, it seems likely that this transcriptional response ultimately stems from the vast genomic damage induced by loss of pol ζ function. Consistent with this hypothesis, the innate immune system not only can recognize and mount an interferon response to foreign DNA, but also can respond to endogenous DNA that has escaped from the nucleus. In some instances, this response can halt cell growth providing organisms to shut down propagations of virally infected cells and cells with dangerously fragmented genomes.

Mammalian cells have a host of cytosolic nucleic acid sensors that patrol the cytosol for DNA. One of these, cGAS, is increasingly recognized to be of paramount importance in the induction of an interferon response to both exogenous and endogenous cytosolic DNA [18]. When cGAS binds to double stranded DNA, it activates its enzymatic activity and results in the production of cGAMP, a cyclic dinucleotide. cGAMP binds to the STING receptor on the membrane of endoplasmic reticulum, resulting in activation of kinases including TBK1 which can in turn phosphorylate and activate IRF3, a central transcription factor in the interferon response.

Given that cGAS-STING axis has been implicated specifically in responding to endogenous DNA damage and has been correlated with micronuclei formation, we asked whether cGAS-STING promotes the induction of expression of interferon stimulated genes due to loss of function of pol ζ. Consistent with most MEFs having a functional innate immune system, both *Rev3l* KO and HET MEF cell lines expressed both cGAS and STING (Fig 3C). We noted a decrease in STING expression in *Rev3l* KO MEFs, which is consistent with a constitutive activation of the cGAS-STING pathway, as cGAS activation leads to a negative feedback loop resulting in STING degradation [19,20]. Importantly, we detected an increase of IRF3 phosphorylated at S888 (corresponding to S396 in humans) indicative of IRF3 activation in *Rev3l* KO MEFs [21,22] (Fig 3D).

This led us to investigate if cGAS-STING drives expression of interferon-stimulated genes upon loss of pol ζ. Knockdown of either cGAS or STING significantly reduced the mRNA expression of selected interferon stimulated genes as well as the protein levels (Fig 4A-D). In addition, depletion of cGAS or STING in *Rev3l* KO MEFs markedly reduced S888 phosphorylation of IRF3 (Fig 4E). Together this indicates that disruption of pol ζ function promotes activation of an innate immune response driven by the cGAS-STING axis.

**Figure 4:**
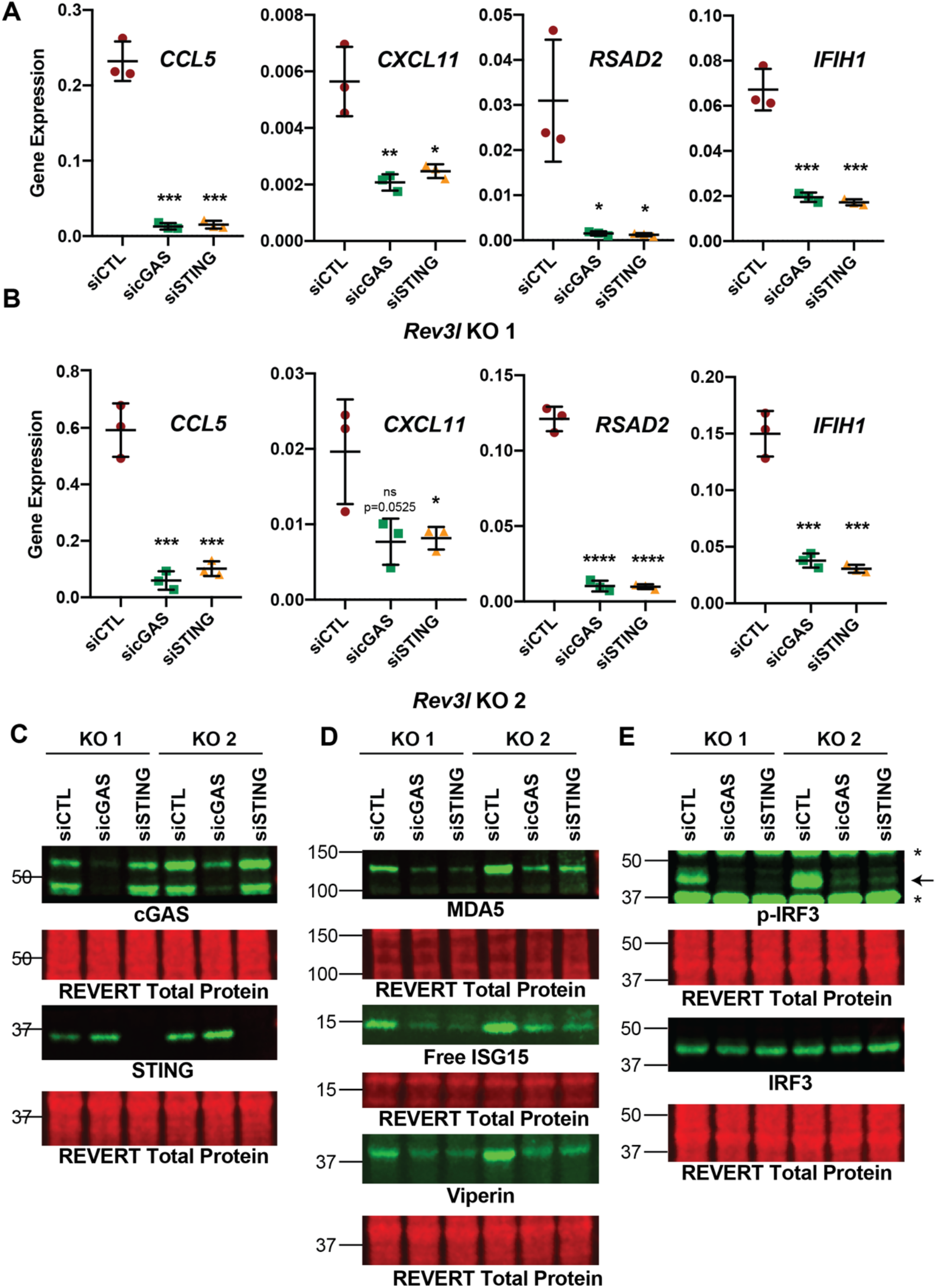
The cGAS-STING axis promotes expression of interferon stimulated genes due to loss of pol ζ function. A) Knockdown of cGAS or STING reduces mRNA expression of *CCL5, CXCL11, RSAD2* (which encodes Viperin protein), *IFIH1* (which encodes MDA5) as detected by qRT-PCR in *Rev3l* KO 1. Gene expression (2^−ΔCt^) of selected interferon stimulated genes normalized to HPRT detected by qRT-PCR. Error bars represent standard deviation. Unpaired student t-test, * = p < 0.05, ** = p < 0.01, *** = p < 0.001, and **** = p < 0.0001. B) Same as in A except with *Rev3l* KO 2. C) Efficient knockdown of cGAS or STING protein levels. D) MDA5, ISG15, and Viperin protein levels decrease with cGAS and STING knockdown. E) Phosphorylation of S888 in mouse (analogous to the human S396) of IRF3 in *Rev3l* KO MEFs decrease with knockdown of cGAS and STING. Equal protein loading was monitored by REVERT total protein stain (**Fig S4**).

**Figure 5:**
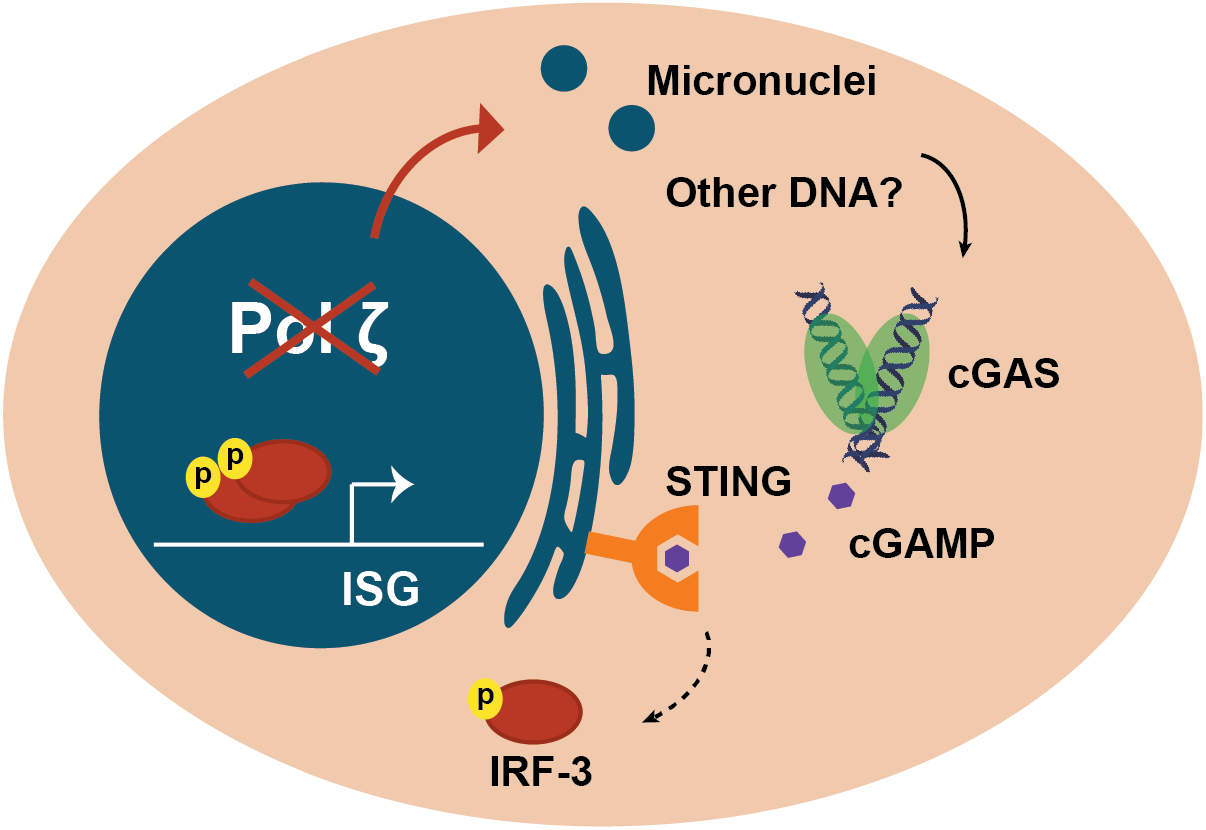
Model for disruption of pol ζ triggering an innate immune response. Loss of pol ζ induces genomic damage that results in accumulation of forms of cytosolic DNA, including micronuclei at a minimum. This results in DNA binding of cGAS and activation of STING which indirectly promotes phosphorylation and activation of IRF-3. This results in expression of interferon stimulated genes (ISG).

### An innate immune response cause by loss of pol ζ function

Pol ζ stands apart from the other translesion polymerases in that it is required for mammalian development and proliferation of primary cells. Now we can add that in addition to activating p53-dependent responses, disruption of pol ζ function invokes a prominent innate immune response promoted by the cGAS-STING pathway.

It is remarkable that disruption of an enzyme commonly thought of as a specialized transleion synthesis polymerase can lead to a constitutive innate immune response. Recently, cells with loss of function of key DNA repair enzymes, RNaseH2, BRCA2, and BLM have been shown to have an elevated cGAS-STING response that correlates with an increase in micronuclei that colocalize with cGAS [23–26]. There are several sources of DNA damage that may give rise to a sustained response including cytosolic mitochondrial DNA [27] and cytosolic DNA arisen from stalled and processed replication forks [28]. DNA stress may continually arise from likely collapse of DNA replication forks in the absence of pol ζ, which could promote formation of micronuclei and also release small fragments of DNA. Further, some nuclear genes including pol ζ control mitochondrial DNA integrity, and there is evidence that mitochondrial function is compromised without pol ζ [29]. It remains to be seen whether micronuclei are the primary source of interferon signalling in cells lacking pol ζ, or whether they are more of an indicator of DNA degradation.

An interferon response can result in shutting down cell growth. Specifically, cGAS has been tied to promoting senescence in primary cells [30,31]. Our experiments were performed in T-antigen immortalized MEFs. In addition to blunting p53 activity, large T-antigen has been implicated in impairing an interferon response to nucleic acids [32]. Loss of pol ζ would likely induce an innate immune response of even greater magnitude in primary MEFs. Primary MEFs lacking pol ζ only make it approximately two cell divisions before cell growth completely halts, which is accompanied by an increase in senescent cells [11]. An essential function of pol ζ is also evident from the failure of *Rev3l*-defective embryos to develop, and the from the inability of *Rev3l*-defective primary keratinocytes to proliferate in a mouse model [2,10,11]. It is possible that cGAS-STING drives this severe growth arrest in primary cells due to loss of pol ζ function.

In addition to widening our understanding of the lengths cell go to protect themselves from the genomic damage induced by impairment of pol ζ, these results could have impact on translational approaches. A potential approach, suggested by experiments in laboratory settings, has been to disrupt pol ζ function to enhance chemotherapeutic effectiveness [33–35]. For example, an inhibitor that impairs the interaction of pol ζ with the master regulator REV1 has been developed that sensitizes cancer cells and xenograft tumors to cisplatin treatment [36]. Our work suggests that such inhibitors might also induce an interferon response. This approach would have multiple advantages for therapy by enhancing DNA damage sensitivity, limiting induced mutations, and potentially enhancing a cytotoxic immune response on targeted cells.

## MATERIALS and METHODS

### Cell Lines

The immortalized *Rev3l* knockout (KO) and heterozygous (HET) mouse embryonic fibroblast (MEF) parental cell lines in this study were as described [11]. In brief, the *Rev3l* KO MEFs were generated by T-antigen immortalization of primary MEFs isolated from mouse embryos with one null *Rev3l* and one floxed allele, followed by ex vivo Ad-Cre deletion of the floxed allele, and clonal selection to ensure a homogenous population with the genotype *Rev3l*^−/Δ^. The *Rev3l* HET MEFs were isolated in the same manner expect starting MEFs from embryos with one wild-type and one floxed allele, resulting in MEFs with the genotype *Rev3l*^+/Δ^. The floxed allele was generated by placing loxP sites flanking two conserved exons, corresponding to residues 2776-2860, that contain the three conserved catalytic aspartate residues of REV3L [6]. The null allele replaced these two exons with a lacZ-neo^R^ cassette [5]. In this study two sets of clones were analyzed. Cells were grown in DMEM (Sigma #5796), 10% Fetal Bovine Serum (FBS) and 1 X penicillin/streptomycin (Gibco #15140122).

### Generation of *TR4-2* expressing *Rev3l* KO MEFs

The TR4-2 construct, a gift from Dr. Wei Yang [15], was cloned into the pCDH backbone with an N-terminal Flag-HA tag as the same previously described for the full length *Rev3l* cDNA [37]. The pCDH-Flag-HA-TR4-2 or pCDH-Flag-HA empty vector was stably inserted into the MEF cell lines using lentiviral infection as previously described [17]. Three single clones were isolated for analysis and continually grown in 1 μg/mL of puromycin to ensure stable integration of the construct.

### RNA isolation

RNA was isolated from 1.5 × 10^6^ (Fig 2, RNA seq) 2.5 × 10^5^ cells (Fig 3, gene expression) or 1 × 10^6^ cells (Fig 4, gene expression), using the RNeasy Kit (Qiagen # 74104) following the manufacturer’s instructions including the on-column DNase I digestion (Qiagen #79254).

### Genome-wide mRNA sequencing

#### Library preparation and sequencing

mRNA libraries were prepared using the Illumina TruSeq Stranded mRNA kit following manufacturer’s instructions and 75 base paired end sequencing was run on the Illumina HiSeq 3000.

#### Sequence mapping and identifying differentially expressed genes

The reads were mapped to the mouse genome (mm10) using tophat V2.0.10 and bowtie V2.1.0. Differentially expressed genes which were defined as > |0.5| fold change and < 0.05 FDR were determined using limma_3.20.4, htseq-count V0.6.0 and edgeR_3.8.6. More stringent cut-offs for fold change were used for various analyses as described below.

### Analysis of differentially expressed genes

For gene ontology analysis, genes upregulated more than 4-fold with an FDR < 0.05 were entered into DAVID 6.8 on 02/22/20 and the top 10 GOTERMS_BP_Direct were plotted based on −log(p value) [38,39]. For upstream regulator analysis, differential expressed gene both upregulated and downregulated more than 4 fold with an FDR < 0.05. were entered into IPA **IPA** (**QIAGEN** Inc., https://www.qiagenbioinformatics.com/products/ingenuity-pathway-analysis). Given that upregulated genes are overrepresented in the Rev3l KO differentially expressed genes, this results in an expected bias for upstream regulators that also predominately result in the upregulation of targeted genes which we see in our data set. Since, bias-corrected z-scores are not reported for upstream regulators with a |bias term| > 0.5, we’ve reported the uncorrected z-score here. Given that the majority of differentially expressed genes are upregulated in our dataset, bias is to be expected for transcription regulators that primarily induce expression of genes.

### Gene expression analysis

High Capacity cDNA Reverse Transcription (Applied Biosciences #4368814) was used to prepare cDNA from 1000 ng of total RNA from each sample. qPCR was run iTaq Universal SYBR Green Supermix (Biorad #1725121) on the Applied Biosystems 7900HT Fast Real-Time PCR System. Gene expression of calculated using the 2^−ΔCt^ method normalizing to the HRPT gene. The following primers (5’ to 3’) were used for the mouse target genes:

HRPT forward: CTGGTGAAAAGGACCTCTCG
HRPT reverse: CAAGGGCATATCCAACAACA
CCL5 forward: ACGTCAAGGAGTATTTCTACAC
CCL5 reverse: GATGTATTCTTGAACCCACT
CXCL11 forward: AGGAAGGTCACAGCCATAGC
CXCL11 reverse: CGATCTCTGCCATTTTGACG
RSAD2 forward: ATAGTGAGCAATGGCAGCCT
RSAD2 reverse: AACCTGCTCATCGAAGCTGT
IFIH1 forward: CGGAAGTTGGAGTCAAAGC
IFIH1 reverse: TTTGTTCAGTCTGAGTCATGG
ISG15 forward: CTAGAGCTAGAGCCTGCAG
ISG15 reverse: AGTTAGTCACGGACACCAG
IRF7 forward: CAATTCAGGGGATCCAGTTG
IRF7 reverse: AGCATTGCTGAGGCTCACTT

### Immunoblotting

Cell were lysed (3 million cells /100 μL) in lysis buffer (Tris-HCl: 50 mM, pH 7.5, NaCl 250 mM, EDTA 1 mM, Triton X-100 0.1%, 1 X Protease/Phosphatase Inhibitor Cocktail CST #5872) for 30 min on ice with mixing every 10 min. Debris was pelleted by centrifugation at 15,000 x *g* for 20 min at 4°C. Protein amounts were quantified using Biorad Protein Assay (Biorad **#**500-0006) and a bovine serum albumin standard curve (Biorad **#**500-0007) following manufacturer’s instructions. The samples were denatured using 4 x loading buffer (LI-COR #928-40004). 25 μg of protein / well and Precision Plus Protein All Blue Standards (Biorad #161-0373) were run on 4-20% polyacrylamide gels (Biorad #4561096) in 1 x Tris/Glycine/SDS buffer (Biorad #161-0772). Protein was transferred to Immobilon-FL PVDF Membrane (Millipore #IPFL00010) in 1 x Tris/Glycine buffer (Biorad #161-0772) 20% methanol. After transfer, membranes were dried. Total protein was measured using REVERT total protein stain kit (LI-COR #926-11010) following the manufacturer’s instructions. Membranes were blocked for 1 h in 0.5X Odyssey Blocking Buffer (OBB, LI-COR # 927-50000) in Tris Buffered Saline (TBS) and then incubated in primary antibody overnight at 4°C. The primary antibodies were used at the following dilutions in 0.5XOBB/TBS/0.2% Tween-20. Rabbit anti-HA-Tag (C29F4) (CST # 3724, 1:1000), Rabbit anti-cGAS (Mouse specific) (D3O8O) (CST # 31659, 1:1000), rabbit anti-STING (D2P2F) (CST **#** 13647, 1:1000), rabbit anti-MDA-5 (D74E4) (CST **#** 5321, 1:1000), rabbit anti-ISG15 (CST **#** 2743, 1:500), rabbit anti-Phospho-IRF-3 (Ser396) (4D4G) (CST #4947, 1:1000), rabbit anti-IRF-3 (D83B9) (CST #4302, 1:1000), mouse anti-viperin (Millipore #MAB106, 1:250), rabbit anti-STAT1 (D1K9Y) (CST: #14994, 1:1,000), rabbit anti-TBK1 (D1B4) (CST: #3504, 1:1,000), and rabbit anti-Phospho-TBK1 (S172) (D52C2) (CST: #5483, 1:1,000).

After primary incubation membranes were washed three times in TBS/0.1% Tween-20, and incubated for 1 h in secondary antibody either goat anti-Rabbit 800CW (LI-COR #926-32211) or goat anti-mouse 800CW (Licor 827-08364) diluted 1:20,000 in 0.5XOBB /TBS /0.2% Tween-20/0.01% SDS. After primary incubation membranes were washed three times in TBS/0.1% Tween-20, rinsed with TBS, then dried and imaged on the LI-COR Odyssey FC.

### Knockdown of cGAS and STING protein levels

600,000 cells were seeded in 10 cm plates. The next day, cells were transfected with 1 nM of the appropriated siRNA using Lipofectamine RNAiMAX (ThermoFisher #13778150) following the manufacturer’s protocols. The following dicer-substrate short interfering RNAs were used: siCTL (IDT: Negative Control DsiRNA # 51-01-14-03), sicGAS (IDT: DsiRNA Duplex mm.Ri.Mb21d1.13.1), and siSTING (IDT: DsiRNA Duplex mmRi.Tmem173.13.2). After 48 h, cells were harvested in paired pellets for RNA and protein analysis and flash frozen in liquid nitrogen.

### Cisplatin sensitivity

10,000 cells were seeded in triplicate in 96 well plates. The following day cells were treated with the appropriate concentration of cisplatin or vehicle control. After 48 h, the relative survival of the cells was calculated by using the ATPlite assay (Perkin Elmer #606016943) following the manufacturer’s instructions.

### Micronuclei frequency

20,000 cells per chamber were seeded on four-chamber slides. After 48 h, cells were fixed in 100% methanol and slides were stained with DAPI. The slides were mounted and imaged. Micronuclei were counted as discrete units distinct from the nucleus.

### Statistics

Unpaired student t-tests were run on qPCR and micronuclei frequency results using Prism 8.

## ACKNOWLEDGEMENTS

We appreciate helpful discussions and advice from MD Anderson Cancer Center colleagues. We thank Drs. Yue Lu and Bin Liu for invaluable assistance with expression data analysis. We thank Winnie Cheng, for assistance with the micronuclei experiments. Studies of pol ζ in our laboratory funded by National Institutes of Health grant CA193124, Department of Defense grant W81XWH-17-10239, and the Grady F. Saunders Ph.D. Distinguished Research Professorship to RDW. SKM was supported by a CPRIT Research Training Grant award (RP170067). The Next Generation Sequencing Core was supported by CPRIT grants RP120348 and RP170002.

## AUTHOR CONTRIBUTIONS

SKM designed and performed the experiments, led data interpretation, and drafted the manuscript. JT established the complemented cell line pairs and assisted with the manuscript. RDW assisted with experimental design, data interpretation and writing.

## SUPPLEMENTARY FIGURES

**Figure S1:**
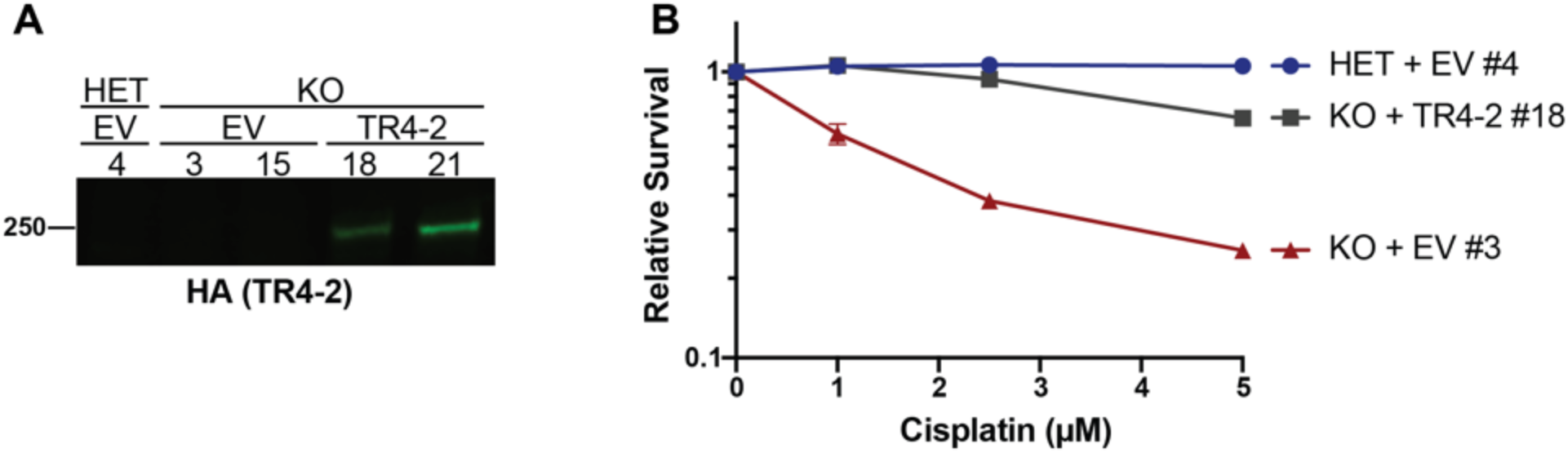
Shortened REV3L construct rescues phenotypes of pol ζ disruption in an additional set of clones. A) Stable expression of TR4-2 with an N-terminal Flag-HA tag in *Rev3l* KO clones as detected by HA immunoblot. See Fig S1 REVERT total protein loading control. B) Stable expression of TR4-2 in *Rev3l* KO MEF clones reverses hypersensitivity to cisplatin. MEFs were exposed to the indicated cisplatin concentrations for 48 hr and relative survival was quantified with the ATPlite assay.

**Figure S2:**
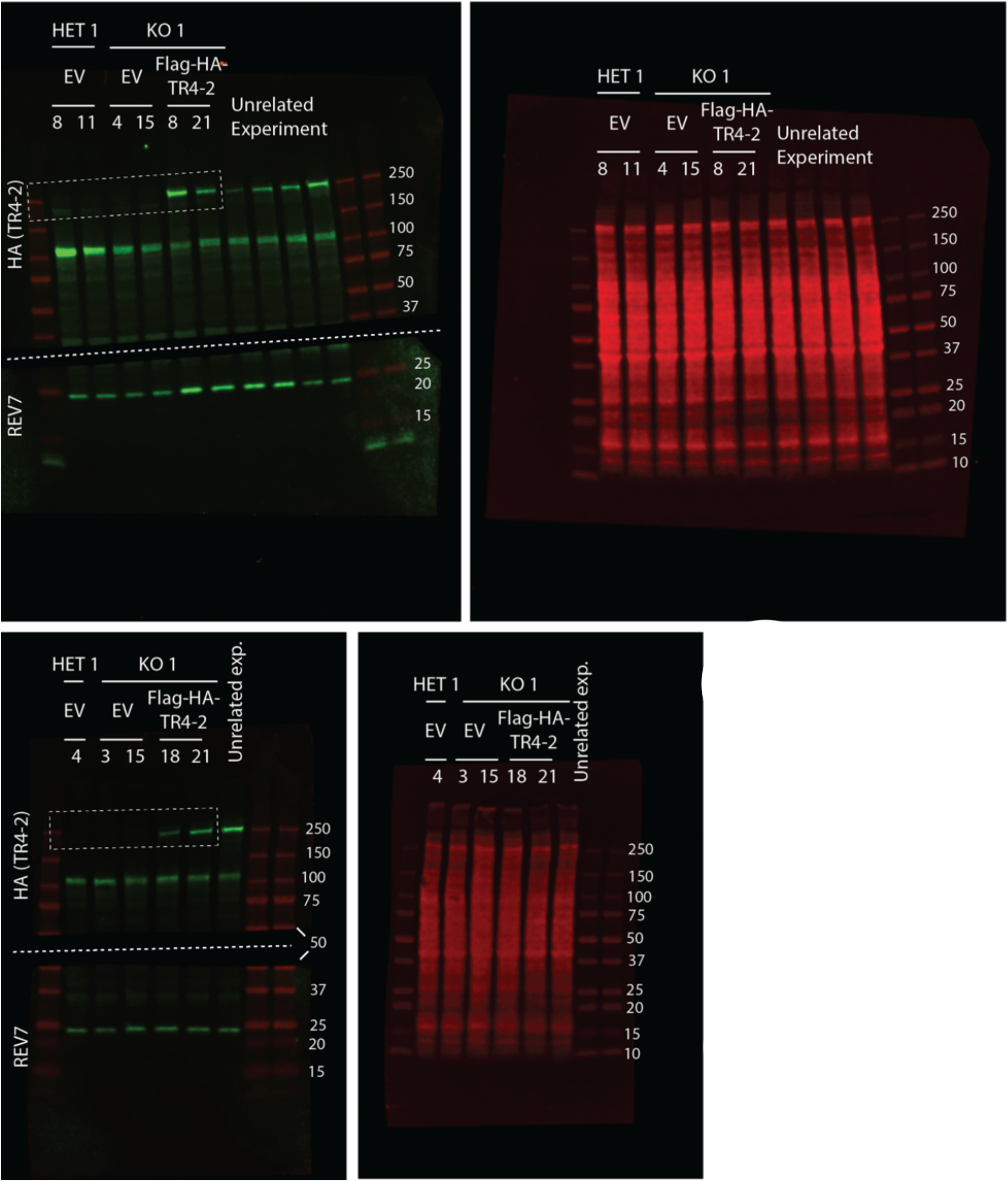
Full immunoblots for Figure 1 and S1. REVERT Total Protein Stain was used as a loading control for each membrane.

**Figure S3:**
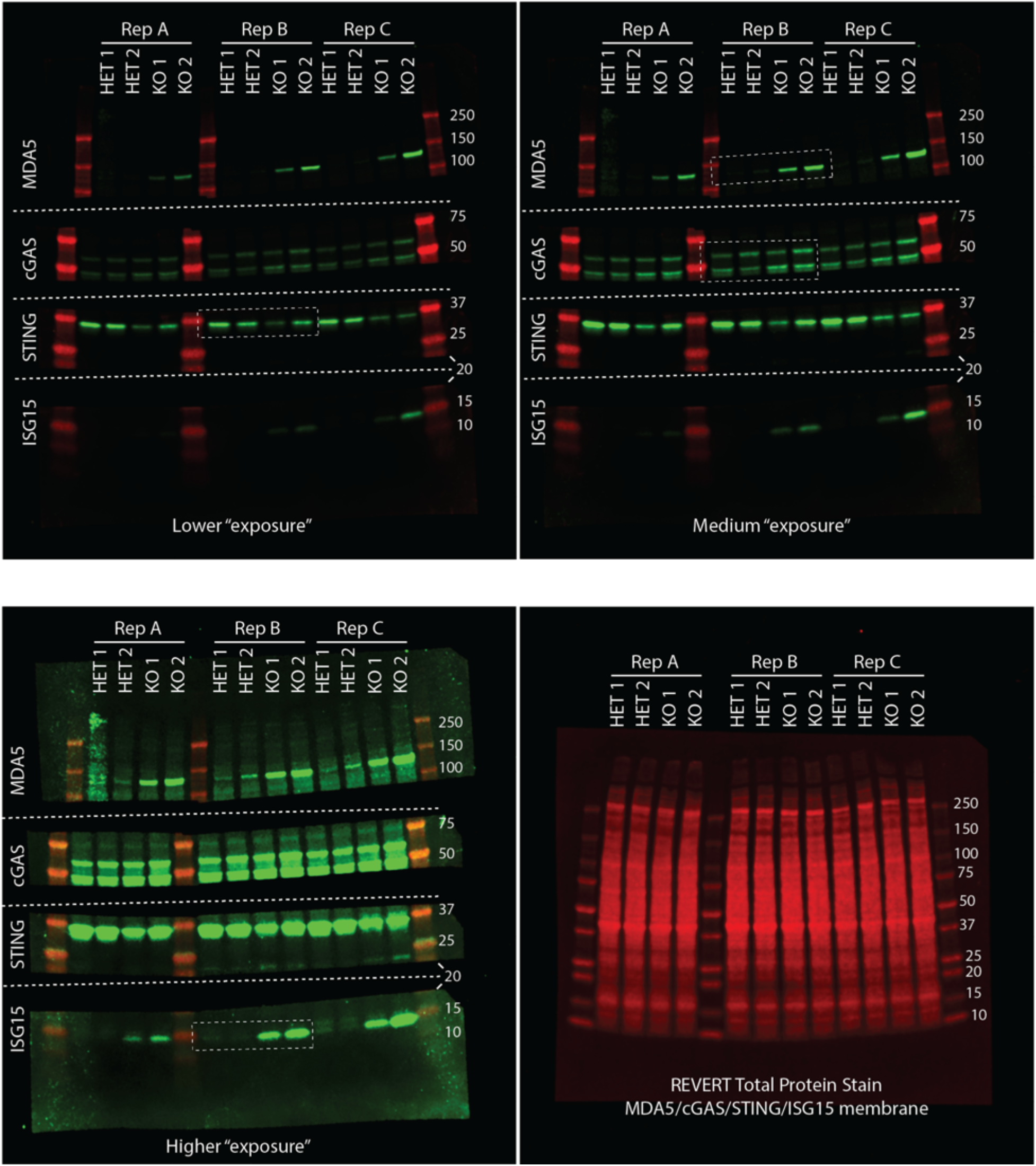
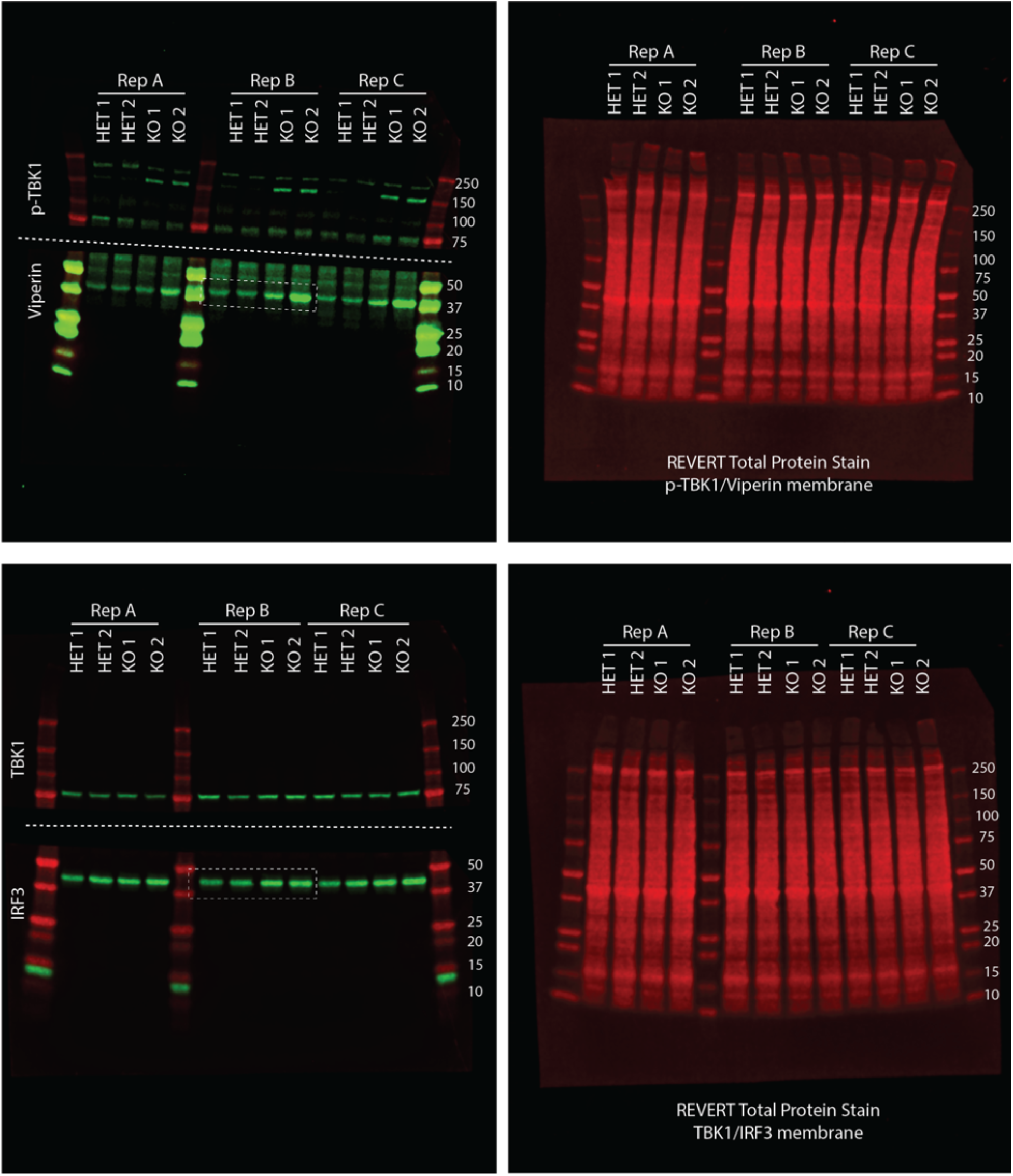
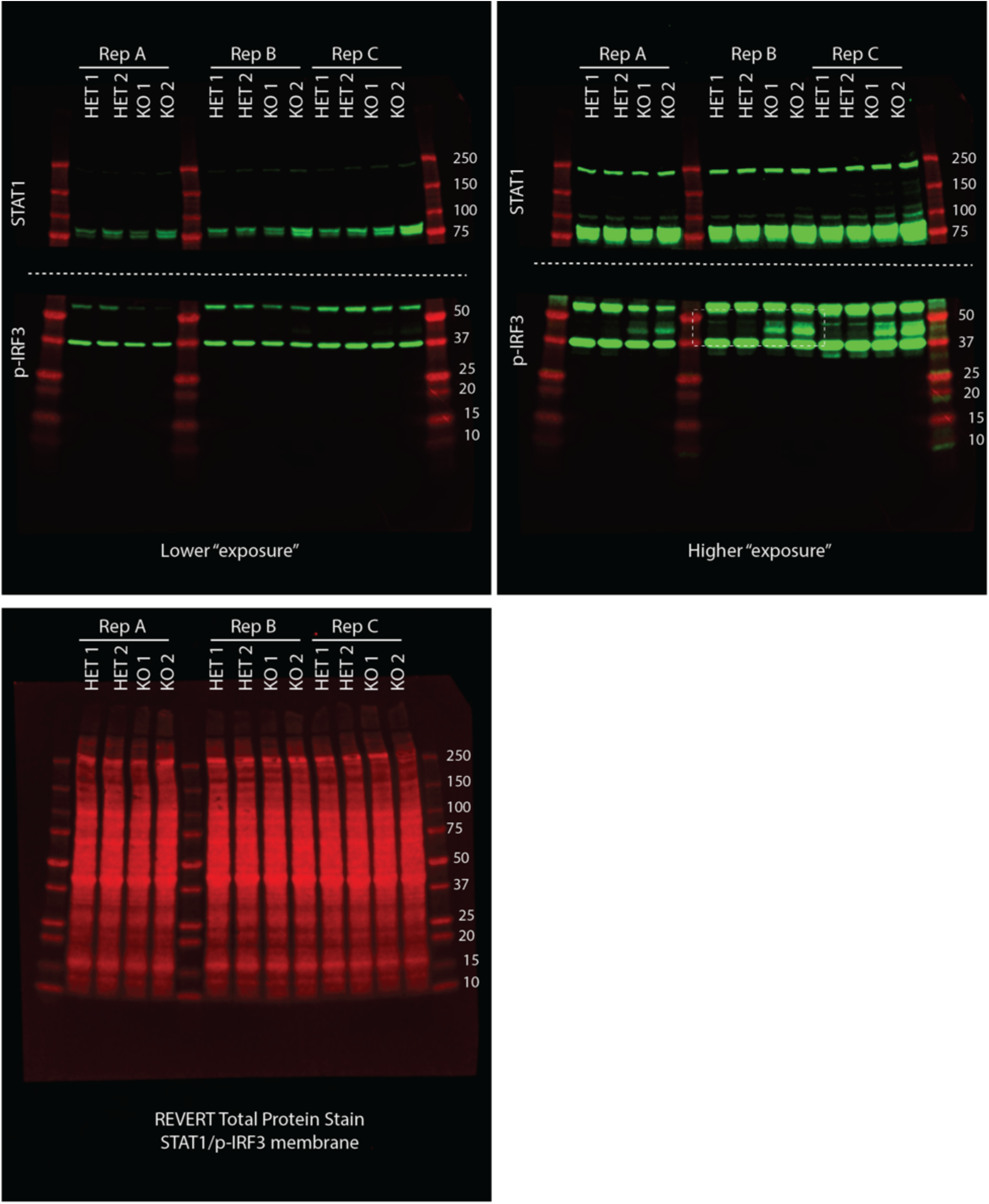
Full immunoblots for Figure 3. Three independent experiments were run on the same gels. Replicate B is shown in figure 3 as indicated by the white boxes. REVERT Total Protein Stain was used as a loading control for each membrane. TBK1, phosph-172-TBK1, and STING were also probed.

**Figure S4:**
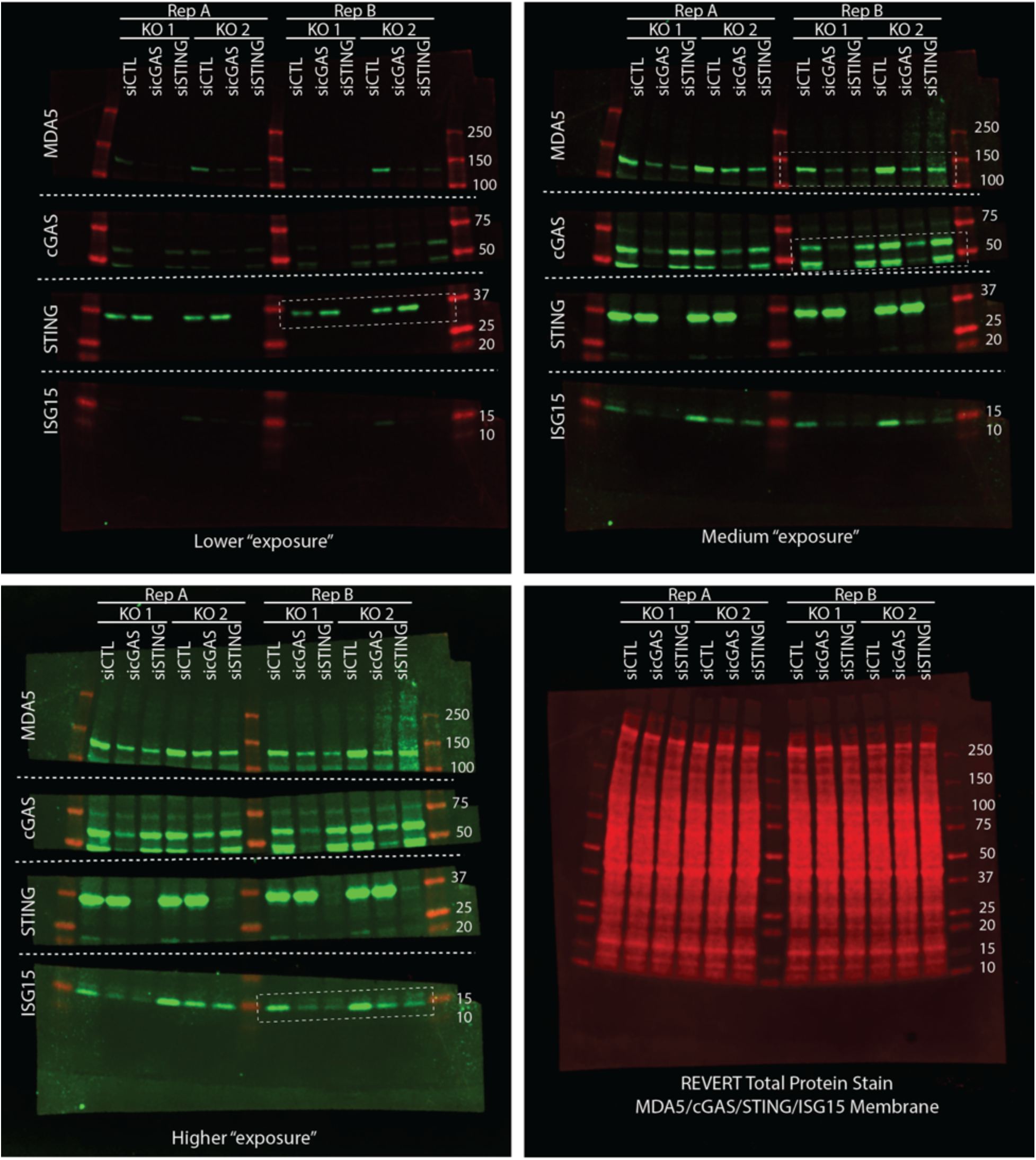
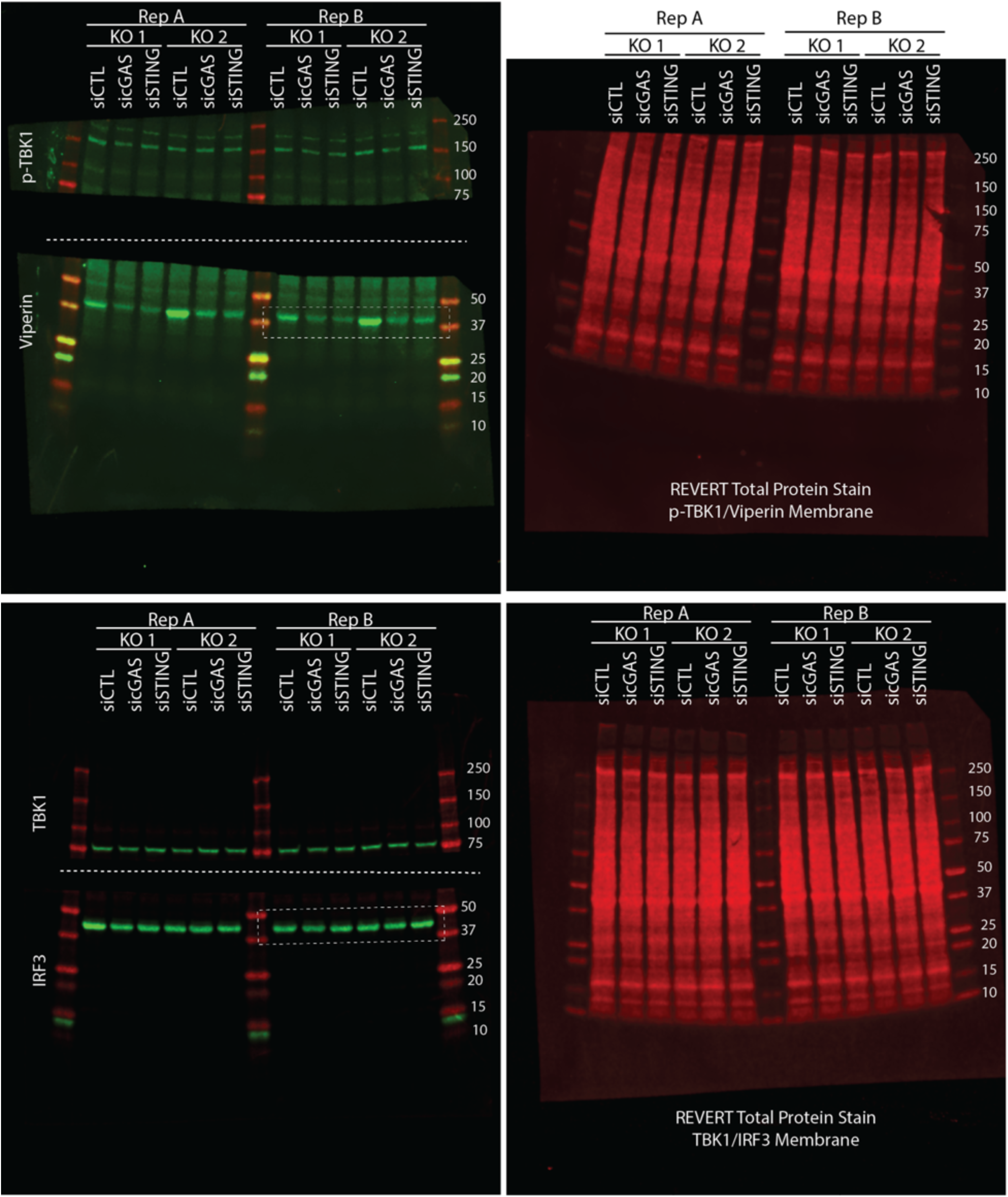
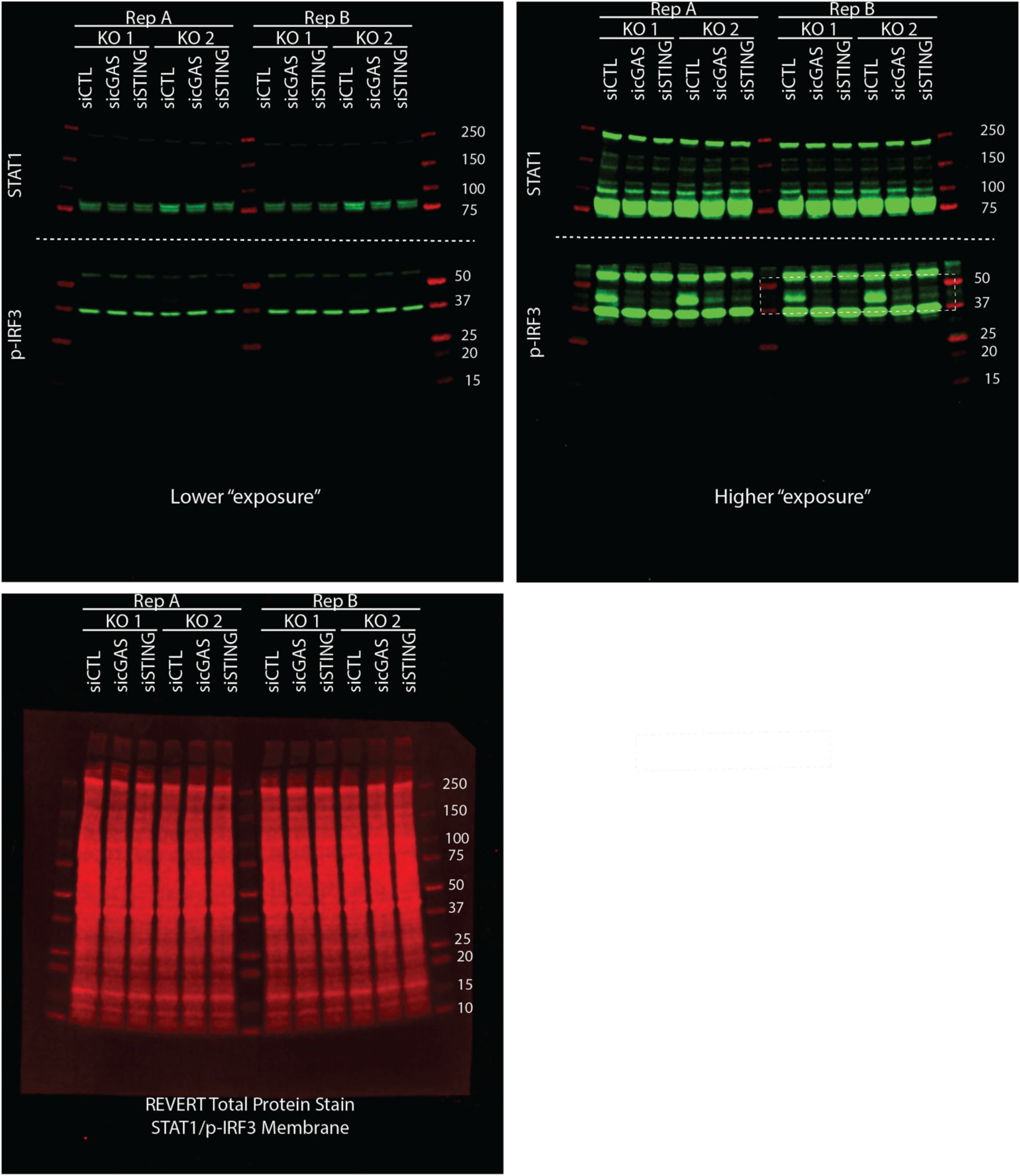
Full immunoblots for Figure 4. Two independent experiments were run on the same gels. Replicate B is shown in figure 4 as indicated by the white boxes. REVERT Total Protein Stain was used as a loading control for each membrane. TBK1, phosph-172-TBK1, and STING were also probed.

